# Dynamic genomic constraints reveal fitness trade-offs underlying bacterial resistance evolution

**DOI:** 10.1101/2025.08.20.671315

**Authors:** Lucy Dillon, James O. McInerney, Christopher J. Creevey

## Abstract

Antimicrobial resistance (AMR) is often modelled as the accumulation of resistance genes leading to multidrug resistance (MDR). We show that gene co-occurrence patterns in two opportunistic pathogens are consistent with fitness trade-offs that constrain which combinations of resistance mechanisms coexist. We applied a combined pangenomic and machine-learning analysis to 9,584 Escherichia coli genomes (99.2% phylogroup B2) and 7,057 Pseudomonas aeruginosa genomes. In E. coli, we identified eight cases of mutually exclusive gene pairs that independently predicted the same MDR phenotype, suggesting alternative routes to resistance whose components are typically not co-inherited. In a separate dataset of 352 strains with paired minimum inhibitory concentration (MIC) data, these dissociated combinations co-occurred more often in resistant than susceptible strains, consistent with the constraints being conditional on antibiotic selection. 33 gene pairs showed opposing association patterns between the two species, with combinations significantly associated in one species and significantly dissociated in the other (e.g. associated in E. coli and dissociated in P. aeruginosa, or vice versa). This indicates that genomic context modifies the contribution of individual genes to resistance phenotypes, and offers one explanation for the observation that 106 ARGs are present in >95% of strains yet do not predict resistance phenotype on their own. The findings are consistent with resistance evolution being shaped by fitness trade-offs and suggest that the dissociation patterns we identify could be targets for follow-up experimental work on resistance-associated fitness costs.

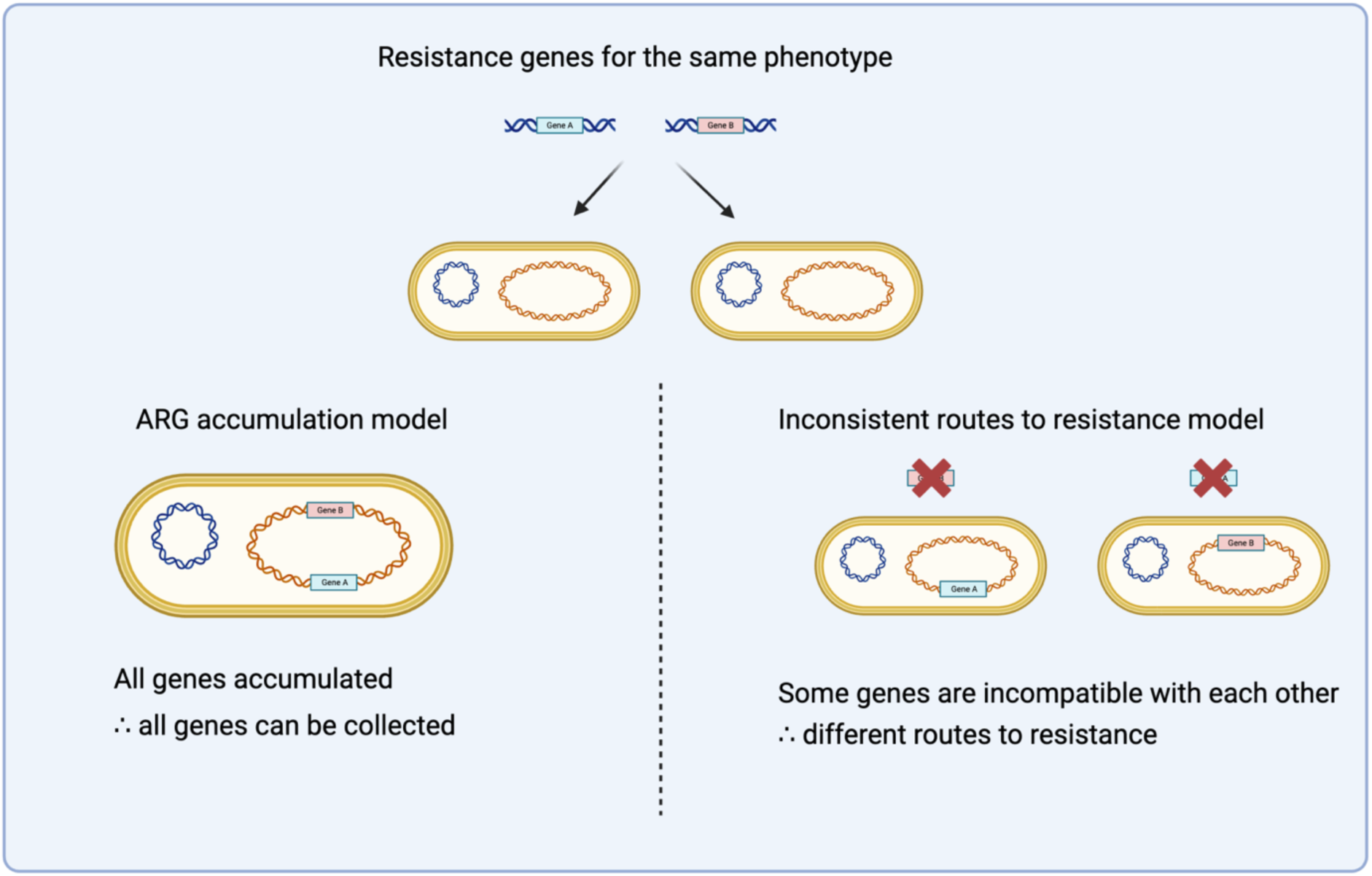

**IMPACT STATEMENT:** Multidrug resistance threatens global health; however, resistance acquisition remains poorly understood. We show that genomes cannot simply accumulate all available resistance mechanisms. Instead, species-specific grammatical rules render certain gene combinations typically incompatible. Analysing 9,584 *Escherichia coli* genomes, we identify eight mutually exclusive routes to resistance, demonstrating that resistance is more constrained than previously thought. Strikingly, 33 gene pairs that cooperate in *E. coli* actively disassociate in *Pseudomonas aeruginosa*, revealing that genomic context determines resistance outcomes. This explains why 106 resistance genes present in all strains fail to confer universal resistance, challenging current diagnostics. Our findings transform our understanding of resistance evolution from an unlimited accumulation model to a grammatically constrained pathways model, presenting new therapeutic strategies that exploit evolutionary dead ends.

## INTRODUCTION

The relationship between phenotype and genotype is rarely as straightforward as the presence of a single gene determining the expression of a particular phenotype (1) and this complexity is particularly evident in antimicrobial resistance. Many phenotypes arise from the interplay of multiple genes, which construct complex mechanisms (2). Gene expression also responds to environmental conditions, including stress (3), or exposure to antibiotics, which can introduce various changes to gene expression regulation (e.g. regulation of porins) (4). Understanding these complex relationships is essential for predicting and combating the evolution of multidrug resistance, yet we lack fundamental knowledge of how bacteria balance the fitness costs and benefits of resistance mechanisms. Complicating the issue further, prokaryotic genomes exhibit considerable variation within a species, which can be represented as a pangenome (5). Prokaryotic pangenome variation is not random but shows highly structured associations in gene content, likely driven by natural selection (6). For instance, the presence of any gene is often related to the presence of its functional partners (7), and the wider genomic content can determine the essentiality of a gene to cell viability (8).

How these pangenomic fitness effects relate to an organism’s traits and phenotypes is still unclear, but it follows that the benefit of any gene to a particular phenotype may also be conditioned on the wider genetic environment. This means that the predictability of a trait or phenotype of an organism may depend not only on the presence or absence of specific genes but also on the wider genomic context in which they are found, for example, the link between antimicrobial resistance (AMR) phenotype and antimicrobial resistance gene (ARG) presence might not be straightforward. The complexity of these dependencies is key to understanding how AMR mechanisms are constructed, and this understanding, in turn, offers a possible avenue to combating resistance. The current trajectory of AMR predicts that there will be 10 million deaths annually as a result of AMR infections by the year 2050 (9). Therefore, urgency is required to understand how AMR arises and how this develops into multidrug resistance (MDR).

Fitness costs are commonly associated with AMR mechanisms, and ARGs are often lost when antibiotic exposure is removed (10,11). The fitness cost can vary considerably depending on many factors; for example, different types of AMR mechanisms (10,12), the AMR gene location (i.e. AMR genes on chromosomes on average have a larger fitness cost than AMR genes on plasmids) (12), and, on average, acquired AMR genes are less costly than AMR mutations (13). AMR mechanisms can impose metabolic costs, including changes to essential metabolic pathways (13). The mechanisms behind these observed fitness costs are not clear and are likely to be driven by not only the presence of a gene but also the genomic context and the environment (14).

Our previous work demonstrated that accurate prediction of AMR phenotypes involves more than simply identifying known antibiotic resistance genes (ARGs) and depends on understanding the complex interactions between known ARGs and the network of accessory non-canonical genes that potentiate or ameliorate resistance (2). Genes can be incompatible with each other for various reasons, for example, gene toxicity (15), issues with genome stability (16,17), functional redundancy (18), and changes in gene expression level (19). As an example, multiple plasmids often cause stability issues, such as I-complex plasmids (20). This raises the question of whether genetic incompatibilities and genomic context can also prohibit or constrain how traits and phenotypes can arise.

In this study, we examined the available pangenome of two key opportunistic pathogens, *P. aeruginosa* and *E. coli* (21,22), to deepen our understanding of their relationship with AMR and MDR, and in particular, the evolutionary pathways through which they arrive at these phenotypes. While MDR is often thought to arise thanks to either MDR-specific genes, plasmids or the accumulation of multiple ARGs (23,24), we have previously shown that the broader genomic background is crucial for understanding which routes to resistance are available to which species (2).

We tested two specific hypotheses using a combination of pangenomics and machine learning: first, that bacteria face evolutionary trade-offs that create conditionally alternative pathways to MDR; and second, that these trade-offs can be overcome under strong selective pressure, revealing dynamic rather than static genomic constraints. Using the *E. coli* and *P. aeruginosa* pangenomes as model systems, we identified 106 unique ARGs across both species that were highly conserved (present in >95% of genomes). Importantly, many of these ARGs (40.1%) could target multiple drug classes, providing numerous potential routes to MDR. Supporting our first hypothesis, we discovered putative incompatible evolutionary trajectories to resistance within *E. coli*, demonstrating that certain combinations of resistance mechanisms typically cannot coexist. In line with our second hypothesis, we found that specific ARGs produced opposing effects in *E. coli* versus *P. aeruginosa*, indicating that, in these cases, genomic background fundamentally alters the route to resistance. These findings demonstrate that the evolution of antimicrobial resistance is not completely deterministic but rather can be contingent upon the broader genomic context in which resistance genes operate. It also shows that genomes generally do not simply accumulate every available ARG. This suggests that future efforts to characterise antimicrobial resistance should account for the broader genomic context.

## RESULTS

### Core genome harbours resistance potential that requires genomic context for phenotypic expression

Using the Roary (25) software pipeline, we built pangenome presence-absence matrices for 7,057 *P. aeruginosa* genomes and 9,584 *E. coli* genomes (see Methods for details). The pangenome analysis revealed that both species have an “open” pangenome (Figure S1A, Table S1) (26), consistent with previous studies (27–29).

First, we interrogated the pangenome matrices to understand whether ARGs are typically found in the accessory genome. We predicted the ARGs using resistance gene identifier (RGI) (v5.1.1) (30) which uses the CARD database. While the majority of ARGs (414/466 in *E. coli* and 360/416 in *P. aeruginosa*) were found in the accessory genome (*<*95% of genomes) (Figure 1B), we identified 106 unique defined ARGs that were present in the core or softcore, which we define as being at least 95% of genomes (56 unique genes for *P. aeruginosa* and 51 unique genes for *E. coli*, one gene was in both species making 106 unique genes) (Figure 1A)). Additionally, 43/106 of the unique ARGs found within the softcore/core genome of both species had MDR genotypes (targeted ≥3 drug classes). The ARGs in the softcore or core genome were predominantly efflux pumps (94/106 unique ARGs; Table S2). This is a challenge for resistance prediction, because genes annotated as “resistance genes” are often essential to bacterial physiology. The dominance of efflux pumps is consistent with exaptation of cellular machinery that originally evolved for housekeeping functions such as heavy metal detoxification, metabolite transport, and membrane homeostasis (31) that can be co-opted for antibiotic resistance under specific conditions. ARG presence alone is therefore insufficient to predict resistance phenotypes, since the functional outcome depends on expression levels, regulatory context, and interaction with other genomic elements (31). It is also known that many of the genes in the CARD database do not confer phenotypic resistance alone, such as *acrA* which is a subunit of a multidrug efflux pump (32), but are part of an overall resistance mechanism. Likewise we have previously shown that ARG identification tools have limits in predicting AMR phenotypes, and more sophisticated techniques, such as machine learning, can provide higher accuracy in predicting AMR phenotypes (2). The remaining 11% of the ARGs found in the softcore and core genome had an alternative function (to efflux) such as antibiotic inactivation or antibiotic target alteration (Table S2).

**Figure 1.**
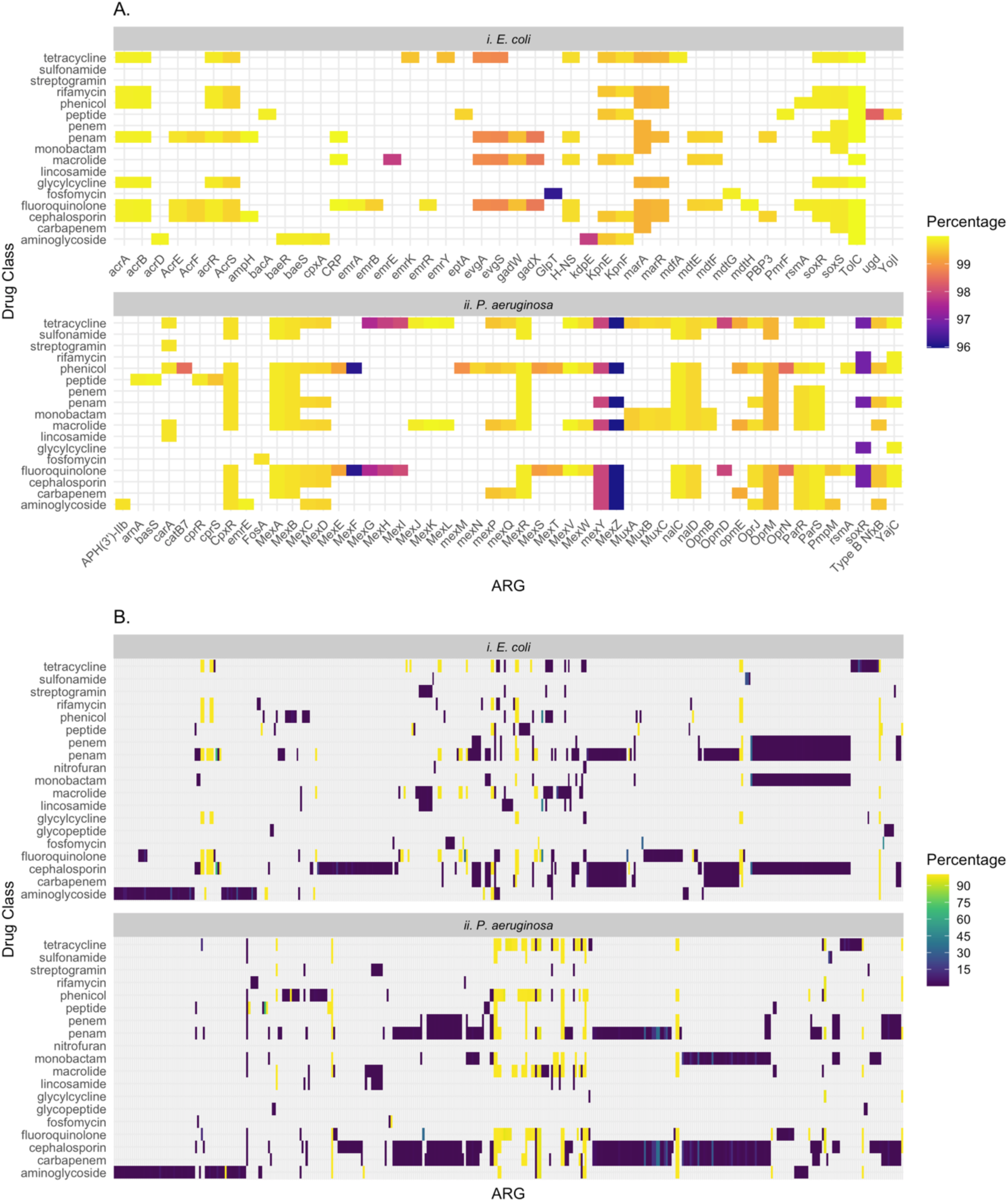
Antimicrobial resistance genes across the pangenome of *E. coli* and *P. aeruginosa.* A. The ARGs present in ≥95% of genomes of *E. coli* (i) and *P. aeruginosa* (ii). The ARGs are shown to correspond to particular drug classes (as documented in the CARD database). The colour bar represents the percentage of the genomes across the pangenome that the gene was found in, specifically highlighting only the softcore and core genomes. A reduced number of drug classes are displayed in the figure for ease of reading. Please see Table S2 for the complete list. Several names have been reduced; please see Table S2 for the exact names in the CARD database. B. The ARG distribution of *E. coli* (i) and *P. aeruginosa* (ii) across the pangenome. The ARG names are not displayed due to the large number of ARGs detected, see Table S2 for full details. Similar to part A, the ARGs correspond to the drug classes, and the colour bar represents the percentage of the genomes across the pangenome that which the gene was found.

### Conditional incompatibilities create fitness-dependent routes to resistance in E. coli

We constructed 55 decision tree models to predict MDR in *E. coli* using an independent dataset of 352 genomes (deliberately withheld from the *E. coli* pangenome analysis) with corresponding laboratory minimum inhibitory concentration (MIC) values for 16 antibiotics (see Materials and Methods) (Figure S1B). The decision tree models provide insight into what genes are key to conferring resistance to antibiotic(s), and they independently validate gene associations from the pangenome analysis (see Figure S2 for an example decision tree). Across the 55 models, mean accuracy was 76.2% but mean precision, recall and F1 were 52.7%, 45.4% and 54.7% respectively (Table S3, Figure S3). Given the four-class MDR phenotype structure (SS, SR, RS, RR) and class imbalance within each comparison, F1 is the more informative summary; several models additionally returned undefined precision, recall and F1 because some phenotype classes received no predictions (Table S3). We therefore use the decision-tree models as a screen for gene-phenotype associations to be cross-checked against Coinfinder, rather than as production-quality predictors.

To understand the routes to MDR in *E. coli*, we used our MDR decision tree models to identify gene families that were inferred to have contributed significantly to a specific AMR phenotype. We found 362 unique eggNOG gene families across all 55 decision tree models. Using these decision tree models, we systematically evaluated the contribution of each gene family to particular AMR phenotypes using a combination of Chi-squared tests, Fisher’s exact tests, and *post hoc* tests. We identified 326 unique gene families (from 519 total instances across the 55 decision trees) that showed significant associations with specific MDR phenotypes (i.e. susceptible to antibiotic A, susceptible to antibiotic B (SS); susceptible to antibiotic A, resistant to antibiotic B (SR); resistant to antibiotic A, susceptible to antibiotic B (RS); resistant to antibiotic A, resistant to antibiotic B (RR)) (Table S4).

To identify putative incompatible pathways to resistance, we combined decision tree analysis with Coinfinder (7) dissociation networks. We specifically tested whether gene pairs that strongly predicted the same resistance phenotype on our decision tree analysis were less likely to co-occur in bacterial genomes than chance expectation. The Coinfinder analysis independently confirmed the predictions from our association analysis for 317 of the gene pairs across the association or dissociation networks. Of these, we uncovered 35 pairs of genes phenotypically important to the same MDR phenotype exhibiting significant genomic disassociation (Figure 2B, Table S4). This suggests that there are putative incompatible evolutionary routes to equivalent resistance phenotypes.

**Figure 2.**
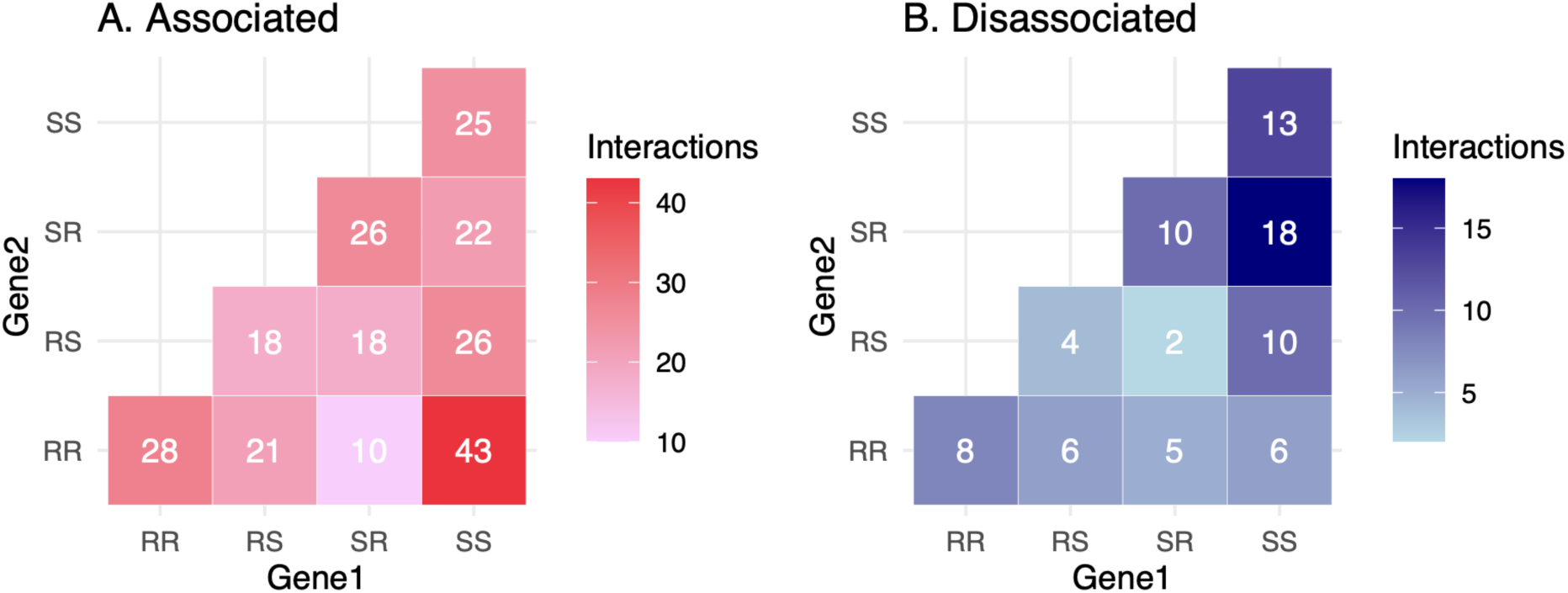
Heatmaps of genes that were statistically linked to a phenotype in the decision trees and then were found to be statistically associated or dissociated with another gene in the Coinfinder networks. Part A shows genes that were associated, and part B shows the disassociations. The phenotype combinations found in the *E. coli* Coinfinder networks are shown.

All combinations of MDR phenotypes were observed in both the association and disassociation networks. For instance, we observed 20 gene families statistically linked to the SS phenotype that were significantly dissociated (in Coinfinder) with gene families statistically linked to the RR phenotype (see Table S4). For example, in the ceftriaxone-ciprofloxacin resistance network the gene family COG3316, which encodes a putative transposon, is significantly linked to dual resistance (RR phenotype) in our decision tree model. On the Coinfinder dissociation network, this gene family showed significant dissociation from the 2AV54 gene family, which itself strongly predicted dual susceptibility (SS phenotype). The fact that gene pairs the decision trees independently link to opposing phenotypes are also genomically dissociated in Coinfinder is an internal consistency check between the two methods.

We also found eight examples of putative mutual exclusion. For example, the two gene families COG3316 and COG3677 are both associated with the RR phenotype (MDR) in the ceftriaxone and ciprofloxacin decision tree model. Nonetheless, the gene families are found to be significantly dissociated in the Coinfinder dissociated network, which indicates that there are putative incompatible routes to MDR phenotypes. The gene families COG3316 and COG3677 are putative transposases (identified using the eggNOG database and searching sequences in the BLAST NT database (33)), this could suggest they are involved in horizontal gene transfer of resistance within *E. coli*.

Figure 2 compares the gene families statistically associated with particular phenotypes in the decision-tree models against the Coinfinder networks in which the same gene-family pairs appear. For example, if two genes are both linked to RR and are found in the Coinfinder dissociated network, this suggests putative incompatible routes to resistance. By contrast, if they are found in the associated network in Coinfinder, they could be part of the same route to resistance. On this basis we identified 27 candidate incompatible routes and 121 gene pairs that may be contributing to the same AMR mechanism (Figure 2). Nevertheless, we cannot explain all instances, such as genes associated with SS-linked genes, since it is difficult to explain associations between genes and the absence of a phenotype that could be beneficial to the organism.

We found that there are Coinfinder associations between genes linked to susceptible AMR phenotypes and the same genes are statistically linked with resistant phenotypes (Figure 2A). We identified three potential explanations for the unexpected associations between susceptibility-linked and resistance-linked genes. First, these associations may represent compensatory relationships where susceptibility-conferring genes are retained to offset the fitness costs of resistance mechanisms. Second, they could indicate epistatic interactions where the phenotypic effect of a gene depends on its genetic partners or the genomic background. In this scenario, a gene promoting susceptibility in one context might be required for resistance machinery to function in another context. This is analogous to the context-dependency of, for example, the *lac* repressor being responsible for prevention of gene expression in the absence of lactose and responsible for expression in the presence of lactose (34). Third, these cases may reflect temporal or conditional dependencies where susceptibility genes are required during non-selective conditions to maintain cellular homeostasis, while resistance genes are activated under antibiotic pressure. Nonetheless, we do not rule out methodological artefacts. The number of isolates used to build our decision trees was fairly low, meaning that the numbers in the exit nodes that were statistically evaluated to assign AMR phenotypes were also quite low. The decision tree models are also built using eggNOG gene families, which are broad gene groups, which could mean that there may be differences within the same eggNOG gene family. Another factor relating to the eggNOG gene families is that they do not take point mutations into consideration, which are documented to be involved in AMR mechanisms (35). Nonetheless, the consistency in the pattern across a broad range of genes suggests biological relevance.

### Co-occurrence of dissociated gene pairs is elevated in resistant strains

The 352 *E. coli* genomes used to build the decision trees with the corresponding laboratory MIC values were used as validation of the Coinfinder dissociated RR labelled COGs, providing confirmation of our computational predictions. When we analysed the eggNOG COGs, for some of the RR dissociated pairs, the dissociated pairs were found together in the same genome for some of the isolates. However, when we analysed the phenotype in conjunction with analysing the dissociation of the COGs, we identified that the dissociation is linked to the AMR phenotype of the isolate (Table S10). We identified that genomes with the RR phenotype had a stronger association between these genes than in genomes with a SS phenotype, in which the genes are mostly dissociated. Therefore, in most cases, these genes are dissociated in “normal” conditions, demonstrating that genomic constraints represent fitness trade-offs rather than absolute rules. Despite this, the RR genes that are typically dissociated may be found together within the same genome because having both RR genes may outweigh the biological burden if exposed to the antibiotic stress. This is consistent with the constraints being conditional rather than absolute, and predicts that strains carrying these combinations would incur a fitness cost in antibiotic-free environments. This could provide insight into what strains may pose potential risks and reveals potential therapeutic targets through exploitation of these evolutionary trade-offs.

### AMR phenotypes are dependent on genomic context

In order to address whether the Coinfinder networks were consistent across the two different pangenomes, we matched the statistically significantly associated gene family pairs found in *E. coli* to the Coinfinder networks of *P. aeruginosa* (Figure S1C). We found 69 matches (gene pair combinations) in the *P. aeruginosa* Coinfinder networks. Not all of the gene family matches were in the same network class for the two species (i.e. associated and dissociate) (Table S6, Table S7). This suggests that similar putative incompatible routes to resistance may exist in both *P. aeruginosa* and *E. coli,* and that these dissociations may be general rather than species-specific. 33 gene-family pairs, however, were assigned to opposing network classes in the two species (Figure 3, Table S7). Figure 3 shows that gene families which are associated in *E. coli* may be dissociated in *P. aeruginosa*. We searched these key gene families for ARGs and were able to identify 15 unique ARGs conferring resistance to 10 different drug classes (Table S8). We detected these ARGs within three gene families, as seen in Figure 3 highlighted in purple. We can see that COG2602 (Beta-lactamase class D) is associated with COG0515 (Carbamoyl phosphate synthase small subunit) and COG0789 (DNA-binding transcriptional regulator, MerR family) is associated with COG1190 (Lysyl-tRNA synthetase, class II) in both *P. aeruginosa* and *E. coli* (Figure 3A). These examples imply that some mechanisms of resistance are conserved across different species (genomic environments). In contrast, COG2271 (a sugar phosphate permease) is associated with COG4388 (Mu-like prophage I protein) in *P. aeruginosa*, but these genes are significantly dissociated in *E. coli*. This is consistent with mechanism components being species-specific, context-dependent, or otherwise plastic with respect to resistance phenotype. These associations allow us to examine how specific genes contribute to resistance phenotypes across different antibiotic combinations.

**Figure 3.**
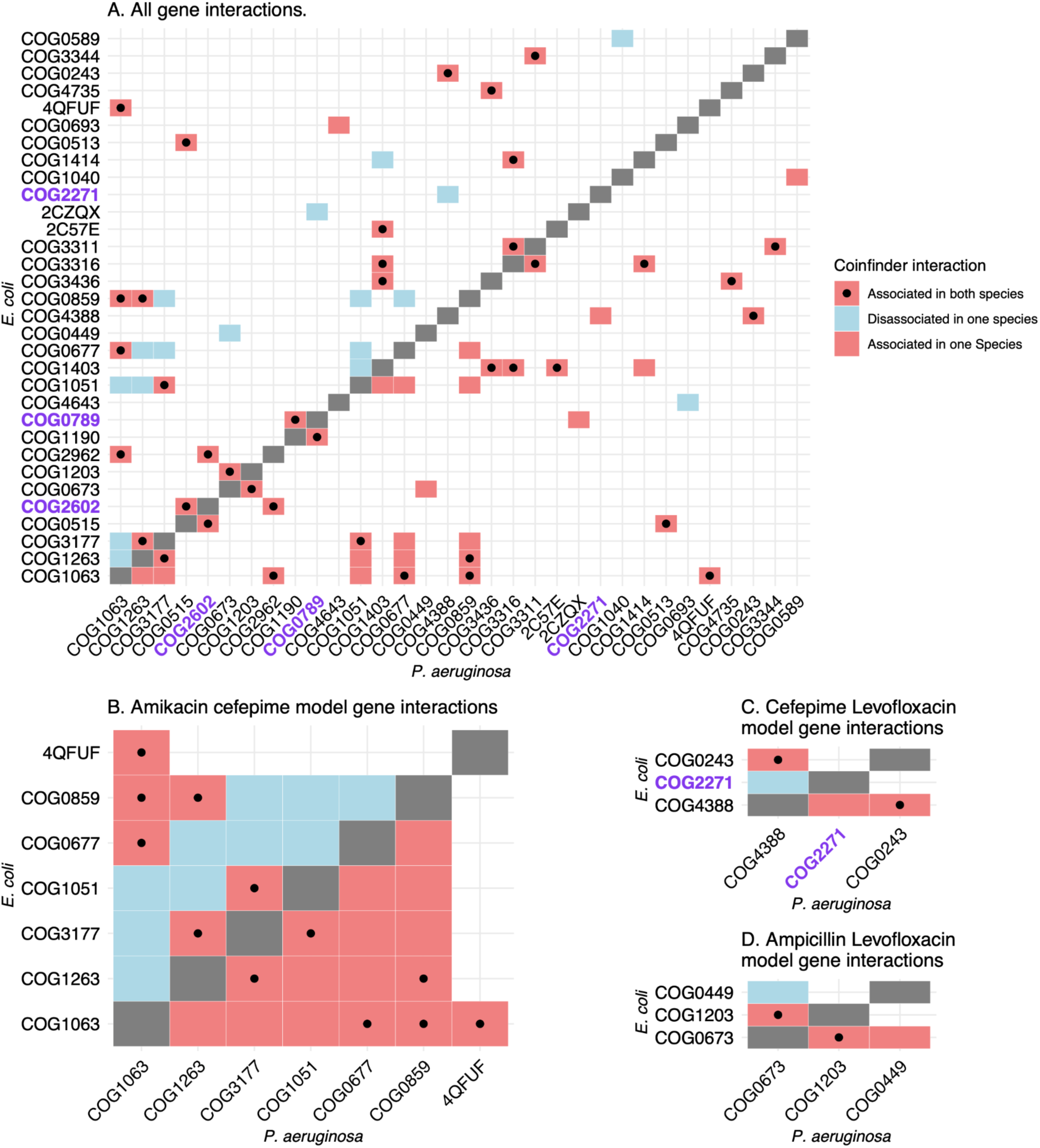
Gene interactions in Coinfinder networks compared in *E. coli* and *P. aeruginosa*. A. All against all gene associations for *P. aeruginosa* and *E. coli*. These genes are statistically linked to MDR phenotypes in the decision tree models (e.g. SS, SR, RS, RR). The plot is split into two triangles split by the grey squares, the lower triangle shows *P. aeruginosa* genes interactions, and the upper triangle shows the *E. coli* gene interactions. The red boxes indicate genes which are associated in Coinfinder, and the blue are dissociated genes in Coinfinder (in the respective Coinfinder species networks). The boxes with a point in the centre represent gene family pairs which share the same association in both species. The three gene families highlighted in purple, when annotated with RGI were identified to have ARGs present. Note -there are no instances of dissociated genes in both species. Parts B-D show genes involved in specific antibiotic models.

The model with the largest number of key genes involved in both *E. coli* and *P. aeruginosa’s* Coinfinder networks was the Amikacin-Cefepime model, with a total of seven genes. 10 of the gene pair combinations manifested contrasting patterns, being dissociated in *E. coli* but associated in *P. aeruginosa* (Figure 3B). COG0859 an ADP-heptose:LPS heptosyltransferase is associated in *P. aeruginosa* for every gene combination available, yet in *E. coli*, three gene families are dissociated (COG3177 (a Fic family signalling protein), COG1051 (ADP-ribose pyrophosphatase), and COG0677 (UDP-N-acetyl-D-mannosaminuronate dehydrogenase)). COG0859 is associated with the phenotype SR (involved in resistance to cefepime, not amikacin). COG3177, COG0677, and COG1051 are all linked to an SS phenotype in *E. coli*. This is most likely why they are dissociated with a gene family that is involved in resistance to cefepime.

In the cefepime-levofloxacin model, three genes are found in both the *E. coli* and *P. aeruginosa* Coinfinder networks. We identified that while COG0243 and COG4388 are associated in both species; COG4388 and COG2271 are associated in *P. aeruginosa* and dissociated in *E. coli* (Figure 3C). Furthermore, we analysed the gene family sequences through the Resistance Gene Identifier (RGI) (36) and identified COG2271 to be linked to AMR, since we identified 15 unique ARG(s) which correspond to 10 drug classes (Table S8).

In the eggNOG database, COG2271 is labelled as being involved in transmembrane transporter activity. Furthermore, the ARGs predicted within COG2271 are involved in efflux transporters (*SoxR*) (37) and importers within the cell (*GlpT*) (38) as defined in the CARD database online (see Table S8 for full ARG names). No function is available for COG4388 in eggNOG, but its phenotype label (SR) matches COG2271, suggesting a role in resistance to levofloxacin but not cefepime. COG1263 is annotated in eggNOG to the phosphotransferase system (PTS); promoter deletion in PTS components has been linked to resistance to beta-lactams and aminoglycosides. Amikacin is an aminoglycoside and cefepime is a beta-lactam (cephalosporin). COG1051 is annotated in eggNOG to GDP-mannose mannosyl hydrolase activity, which has been linked to L-fucose production (39) which has a role in antimicrobial activity. L-fucose is necessary for cell adhesion and biofilm formation (40), processes that are common in opportunistic infections (41,42). Nevertheless, the phenotype of COG1051 is SS. This is most likely dissociated with COG0677, which has an SS phenotype. However, in *P. aeruginosa,* these genes are associated, which suggests that these genes function together in *P. aeruginosa*, but not in *E. coli*.

In the ampicillin-levofloxacin model, three genes are found in both the *E. coli* and *P. aeruginosa* Coinfinder networks. While COG1203 and COG0673 are associated in both species, COG0449 and COG0673 are associated in *P. aeruginosa* but dissociated in *E. coli* (Figure 3D). COG0449 and COG0673 are both statistically linked with the RR phenotype in the ampicillin-levofloxacin decision tree model, however, these genes are genomically dissociated in the *E. coli* Coinfinder network, even though these genes are associated in the *P. aeruginosa* Coinfinder network. COG0449 is involved in glutamine-fructose-6-phosphate transaminase (isomerizing) activity, which is part of the hexosamine pathway (43,44). This gene family is a potential therapeutic target for antimicrobials, which suggests the gene does have a key role in AMR. COG0673 is thought to be involved in inositol 2-dehydrogenase activity (44). Inositol has been shown to influence microbial physiology, such as motility and extracellular matrix phenotypes (45). These features are known to increase the pathogenicity of bacteria (46), which could aid with antimicrobial treatment.

## DISCUSSION

The integration of pangenomics with machine learning has revealed that gene co-occurrence patterns in two opportunistic pathogens are shaped by fitness trade-offs that condition which combinations of resistance components coexist. The MIC-paired dataset is consistent with these incompatibilities being conditional rather than absolute: dissociated gene combinations were tolerated more often in strains where both genes contribute to resistance against the selecting antibiotics. These results are difficult to reconcile with models in which resistance is acquired purely by accumulation of independent ARGs, and instead point to dynamic, species-specific genomic incompatibilities, or a species-specific genomic ‘grammar’ of compatibility rules. Addressing this requires moving beyond gene cataloguing to characterising the wider interactions across the pangenome.

### A “Grammar” of genetic compatibility influences high levels of diversity of MDR mechanisms between *E. coli* and *P. aeruginosa*

In this study, we illustrate the complexity of resistance mechanisms both within *E. coli* and between *E. coli* and *P. aeruginosa*. This complexity challenges the “one gene, one resistance” paradigm that underlies many diagnostic and surveillance approaches (47–51). The presence of 106 ARGs in the core pangenome, with 56.8% targeting multiple drug classes, demonstrates that resistance cannot be reduced to simple presence/absence matrices. If we assume a simplistic correlation between ARG presence and resistance phenotype, this suggests that all 9,584 *E. coli* and 7,057 *P. aeruginosa* isolates have MDR phenotypes.

Understanding this requires investigating genome diversity and the interactions between genes. Our pangenome analysis of *E. coli* and *P. aeruginosa* highlights the extraordinary plasticity of these species. The minimal core genomes, representing less than 1% of both species was notably small, even compared to previous studies on these organisms (27,52–55). This underscores that each isolate represents a unique resistance evolution experiment. Furthermore, the 317 significant gene associations and dissociations show that this plasticity is not random noise, but structured variation that follows rules imposed by functional constraints. While no two species may share the same pangenome, the evolutionary constraints they face are the same, resulting in independent genetic solutions to the same phenotypic problem. These factors support our hypothesis that there is a species-specific genomic ‘grammar’. In this context we define to genomic ‘grammar’ to mean a set of compatibility rules governing which combinations of gene families co-occur within a species, such that the phenotypic contribution of a gene depends on its co-occurring partners and may not transfer to a different genomic background. Minor differences in the genomic ‘grammar’ could influence gene function within species.

### The Paradox of Infinite Solutions vs. Observed Constraints

Our data shows *E. coli* has putative incompatible evolutionary routes to MDR, suggesting organisms generally cannot accumulate all possible mechanisms of resistance.

We have previously shown that there are often multiple routes to resistance to a single antibiotic, and can often be species-specific (2). Here, we have shown that this extends to MDR phenotypes. Our integrated pangenome-machine learning approach identified that not all resistance mechanisms are always compatible within *E. coli* (Figure 2). Additionally, while certain genes exhibit a specific Coinfinder association in *E. coli*, the same genes may show the opposite association in *P. aeruginosa*, indicating that they may not share the same mechanism across species (Figure 3). This highlights the critical role of genomic context in understanding AMR phenotypes.

The importance of genomic context extends beyond individual gene function to entire resistance strategies. As recently emphasised for biosynthetic gene clusters (56), the genomic neighbourhood fundamentally shapes functional outcomes. Our finding that 33 gene pairs show opposite associations between *E coli* and *P. aeruginosa* is consistent with this, suggesting that the same genetic elements can contribute to resistance in one genomic context while being unfavourable in another. This context-dependence means that resistance markers identified in one species may not transfer directly to others, and that species- or even strain-specific understanding of resistance drivers is needed.

The patterns of putative incompatibilities we observe suggest several possible mechanistic explanations. The incompatibility between COG3316 and COG3677 which are both putative transposases that confer MDR phenotypes, may arise from competition for cellular resources required for transposition, or from destabilising effects of multiple active transposable elements (15–17). Similarly, the 94 efflux-related genes we identified in the core genome likely face constraints on simultaneous expression due to membrane space limitations or energetic costs of maintaining multiple ATP-dependent transport systems. The striking difference in gene associations between *E. coli* and *P. aeruginosa*, particularly for genes like COG2602 (β-lactamase) and COG0515 (carbamoyl phosphate synthase), suggests that regulatory architecture unique to each species determines which resistance combinations are viable. These mechanistic constraints, whether metabolic, regulatory, or structural, shape the evolutionary landscape of resistance and explain why bacteria cannot simply accumulate all available resistance mechanisms.

The SS-RR association patterns are unexpected and could reflect compensatory or co-regulatory relationships between susceptibility-associated and resistance-associated genes; the MIC-paired dataset shows that the pattern is structured by phenotype, although technical contributions from broad eggNOG categories and modest per-class sample sizes cannot be fully excluded. Several non-exclusive biological mechanisms could explain these patterns. For example, resistance mechanisms often impose substantial metabolic costs (57). Genes associated with susceptibility phenotypes might encode functions that maintain cellular efficiency, making them essential partners for energy-intensive resistance mechanisms. For example, improved metabolic efficiency (associated with susceptibility in the absence of antibiotics) might be required to support ATP-dependent efflux pumps. Alternatively, these associations might reflect shared regulatory networks where stress response systems co-regulate both resistance mechanisms and susceptibility-associated housekeeping functions. Under antibiotic stress, the entire regulon is activated, but only the resistance components contribute to survival.

The associations could also represent evolutionary “package deals” where genes cannot be separated due to shared promoters, operons, or mobile genetic elements. This would mean bacteria must accept susceptibility-associated genes to gain resistance capabilities, representing another form of our proposed genomic “grammar”.

Although it is possible that technical factors (broad eggNOG categories, limited sample size) may contribute to some associations, the consistency of these patterns suggests biological relevance. Future work using isogenic mutants and expression studies could definitively determine whether these associations reflect functional dependencies or technical artefacts.

### Fitness-dependent constraints create dynamic pathways to resistance

Toxicity (15), genome stability (16,17), functional redundancy (18), and changes in gene expression level (19) can result from incompatibilities between a variety of genes (20). However, little is known about how the incompatibility of ARGs may influence MDR phenotypes. We identified 8 instances in *E. coli* where the genes forming a route to an MDR phenotype were putatively incompatible with genes that formed a different route to the same MDR phenotype. This suggests that different pathways to resistance may be putatively incompatible. These gene disassociations indicate that specific constraints and trade-offs shape the presence of certain genes, influencing how particular phenotypes emerge in specific organisms. The putative incompatible routes may also be a result of functional redundancy if an isolate already has a resistance mechanism, it might be a genomic burden to have another resistance mechanism. An example of this from our data is the observed dissociation between COG3316 and COG3677 transposases, despite conferring MDR, may actively conflict with each other. These trade-offs represent evolutionary bottlenecks (for instance, if bacteria cannot maintain all possible resistance mechanisms simultaneously) that could be exploited therapeutically by forcing bacteria towards less favourable evolutionary paths. The MIC-paired dataset is consistent with these constraints being conditional: dissociated gene pairs co-occur more often in resistant strains than in susceptible ones (Table S10), suggesting that the fitness cost of carrying both is tolerated only when antibiotic selection makes both components beneficial. If this interpretation is correct, strains carrying these combinations would be predicted to incur a measurable fitness cost in antibiotic-free environments, a prediction that could be tested experimentally.

### The wider genomic context must be considered in MDR treatment

Many hospitals incorporate whole genome sequencing in the diagnostic stages of diseases (58,59). For example, real-time polymerase chain reaction (PCR) can be used in the clinical setting to identify ARGs in patient samples (60). When using pathogen genomes to predict antibiotic resistance, the accuracy of these predictions depends on accounting for taxonomy and other genes that may influence the phenotype. Our findings have immediate implications for clinical practice. First, diagnostic platforms must move beyond binary resistance gene identification to assess fitness trade-offs. Strains carrying “forbidden” gene combinations may be resistant but evolutionarily unstable, suggesting windows for intervention. Second, our identification of conditional incompatibilities suggests that strategic antibiotic withdrawal could select against resistant strains that carry fitness-costly gene combinations. Third, combination therapies could exploit these trade-offs by forcing bacteria towards evolutionary dead ends where resistance comes at a fitness cost. This emphasises the complexity of resistance mechanisms and the challenges they pose for accurate resistance prediction. To improve AMR phenotype predictions, factors such as taxonomy and key accessory genes influencing resistance should be considered. The lack of applicability of “universal” resistance markers across species may explain cases of treatment failure despite “susceptible” genotypes and false resistance predictions despite ARG presence.

The presence of 106 ARGs in the core genome highlights a particular problem with current AMR surveillance strategies that rely on ARG detection. These 106 genes are likely to have “resistance potential” rather than realised resistance. Their universal presence suggests three compatible possibilities: (1) these genes primarily serve essential housekeeping functions with resistance as a secondary capability activated only under specific conditions, (2) resistance phenotypes require additional accessory genes or regulatory changes to manifest, or (3) current ARG databases conflate genes with resistance potential with those that actively confer resistance. This distinction is important for clinical diagnostics -detecting these core ARGs provides no predictive value for treatment outcomes, yet they may be a reservoir of resistance potential that could be activated through regulatory mutations or in combination with accessory genes.

This observation directly supports our genomic grammar hypothesis – that the presence of resistance-associated genes is necessary but may not be sufficient for resistance phenotypes. The grammatical rules determining when these core genes contribute to resistance versus when they simply have housekeeping functions remain to be worked out, but these rules are likely to involve the broader genomic context we investigate here.

### Methodological innovation enables biological discovery

The biological insights presented here were made possible by the convergence of two complementary approaches. Traditional pangenome analysis alone would have identified gene presence/absence patterns but not their phenotypic relevance. Machine learning alone would have predicted resistance but not the evolutionary constraints underlying them. By integrating Coinfinder’s unsupervised network analysis with supervised decision tree learning across 55 antibiotic combinations, we could both identify statistical associations and validate their phenotypic outcomes, revealing patterns missed by either approach alone. The 317 gene pairs identified by both methods demonstrate that novel biological insights can be observed when we move away from single analytical frameworks to approaches that capture both evolutionary and functional outcomes.

## Conclusion

Our combined pangenome and machine-learning analysis is consistent with bacterial resistance evolution being shaped by conditional fitness constraints, and identifies candidate gene pairs whose dissociation under non-selective conditions could be targets for follow-up experimental work. We identified eight conditionally incompatible routes to MDR in *E. coli*, and the MIC-paired dataset is consistent with bacteria pay fitness costs for maintaining “forbidden” gene combinations under antibiotic pressure. 33 gene pairs show opposite associations between *E. coli* and *P. aeruginosa* is consistent with these constraints are species-specific, challenging the portability of resistance markers across species.

These findings have immediate clinical relevance. Rather than viewing resistance as inevitable accumulation, our work is consistent with resistance evolution being shaped by a balance between survival benefit and fitness cost, which suggests directions for follow-up experimental work. In principle, diagnostic platforms could identify strains carrying combinations expected to incur a fitness cost in antibiotic-free environments, and treatment strategies might exploit fitness trade-offs to select against resistance. The presence of 106 ARGs in core genomes yet variable resistance phenotypes underscores that gene presence alone cannot predict treatment outcomes.

Together, these findings indicate that bacterial resistance evolution is more structured by compatibility constraints than a simple accumulation model would predict, opening avenues of research in the fight against multi-drug resistance.

All computational pipelines and data are freely available (https://github.com/LucyDillon/MDR_pangenome) to enable replication and experimental follow-up.

## MATERIALS AND METHODS

Within this study, we investigated AMR phenotype within a species (*E. coli*) and between species (*P. aeruginosa*) using the genomic context provided by pangenomics and machine learning models.

### Data availability

To ensure full reproducibility of our pangenomic findings, we provide: (1) all genome assemblies analysed (accession lists at https://github.com/LucyDillon/MDR_pangenome/), (2) Roary pangenome matrices for both species and (3) Coinfinder association/dissociation gene pairs are available on OSF, (4) trained decision tree models for all 55 antibiotic combinations (.arff format), (5) RGI annotations mapped to pangenome positions (Table S2) (https://osf.io/95k6z/?view_only=b46e8760a93b4e70a849fe4d2210165f), and (6) complete analysis pipeline including sourmash clustering parameters (https://github.com/LucyDillon/MDR_pangenome).

View supplementary information at this link: https://osf.io/95k6z/?view_only=b46e8760a93b4e70a849fe4d2210165f

### Code availability

Please see https://github.com/LucyDillon/MDR_pangenome for all files and scripts used in this study. This includes scripts to make figures in R (see the “Figures” directory in the GitHub link).

Apply decision tree software: https://github.com/ChrisCreevey/apply_decision_tree

### Data for Analysis

To carry out the pangenome association/disassociation analysis, genomes were downloaded from BV-BRC (61), 37,451 *E. coli* and 7,109 *P. aeruginosa*, respectively. These genomes were selected from BV-BRC using the “complete” or “WGS” genomes and “good quality”, and all genomes were filtered to have less or equal to 500 contigs (to reduce the number of genomes and increase the quality). Genomes used in our previous models (2) were removed from the list (see E_coli_MDR_tree_genome_ids.txt). Using Sourmash (version 4.6.1) (62), we clustered the genomes to remove redundancy and misclassified species, using a Kmer size of 31. Since there was a large amount of *E. coli* genomes, a sub-cluster was chosen; these genomes were more genetically similar to each other and a more computationally manageable number (9,584 genomes) (Figure S4). The advantage of this approach is that a smaller dataset of closely related taxonomy for the pangenome will be more appropriate, resulting in a larger core genome to analyse. To understand the diversity of the subcluster of the *E. coli* genomes, the phylogroup was analysed using EzClermont v0.6.3 (63). We identified that 9504/ 9,584 (99.2%) were assigned to phylogroup B2. Our conclusions about *E. coli* therefore apply specifically to phylogroup B2; generalisation to the wider species will require analysis of a phylogenetically broader sample. This resulted in 9,584 *E. coli* and 7,057 *P. aeruginosa* genomes. We chose both species due to their clinical relevance. To create MDR machine learning models of *E. coli,* we used 352 *E. coli* genomes with corresponding minimum inhibitory concentration (MIC) values to multiple antibiotics from our previous dataset, sourced from BV-BRC (2), to build the MDR models. The MIC values were matched to European Committee on Antimicrobial Susceptibility Testing (EUCAST) breakpoints (64) to determine if isolates were resistant or susceptible. Isolates with intermediate phenotypes were removed from the dataset. The genome IDs for the *E. coli* genomes can be found in E_coli_MDR_tree_genome_ids.txt.

### Pangenome analysis

The genomes were annotated with Prokka (v1.14.5) with default parameters (65). Roary (version 3.13.0) (25) was used to create the pangenome and create the core genome alignment (Roary.sh). Roary was chosen over other pangenome tools due to its reproducibility with prior work published and the computational time for >9,000 genomes was appropriate in comparison to other tools, such as Panaroo, which does not utilise multithreading while processing paralogs hence this means the analysis would take much longer to complete. To create a Newick file (necessary for Coinfinder input), FastTree (v2.1.11) (66) was used with the core gene alignment and an R script remove_zero_branch_length.R was used to edit the tree to remove any zero branch lengths in the tree. Zero length branch lengths were replaced with a small constant (1e-6), as recommended in the Coinfinder GitHub. The Roary output file: gene_presence_absence.csv file was edited to remove any spaces from the index of the file before using Coinfinder (see remove_spaces_help.md for details). Coinfinder (v1.2.1) was then used to find significantly associated and dissociated co-occurring genes across the pangenome using default settings (7). We analysed both associate and dissociate analyses separately on our data (Coinfinder_associate.sh, Coinfinder_disassociate.sh). Gephi was used to visualise the network.gexf files using the Fruchterman Reingold layout (67).

### Functional and ARGs annotation of the pangenome

eggNOG mapper (v2.1.6) was used with default parameters to functionally annotate all the genomes (68) (see Pseudomonas_eggnog.sh and E_coli_eggnog.sh). To match the eggNOG mapper results to the pangenome, the IDs generated using Prokka can be used to link the Roary output (gene presence absence.csv) to the eggNOG file (pipeline commands: eggNOG in pangenome.md). As can be seen in the scripts, once the ID is matched, the number of isolates the gene is present in (in column 4 of gene presence absence.csv) can be returned. This can be matched to the eggNOG COG (gene family). By using the number of isolates, we can determine if the gene is present in the core, softcore, shell or cloud portion of the pangenome. Resistance Gene Identifier (RGI) (v5.1.1) (30) was used with mainly default parameters to identify ARGs across the pangenome, which uses the CARD database (v3.1.1) (69) (see E_coli_RGI.sh and Pseudomonas_RGI.sh). We used a ‘strict’ identification cut-off when running RGI to ensure we reduced the chance of identifying false positives. We also analysed the most common sequence of the protein ORF matched to the RGI gene (column 19 in RGI output file) for ARGs present in >95% of genomes using BLASTp to ensure the sequences were not matching to similar genes that may have a more generic housekeeping function. The ARGs were matched to the pangenome by following the commands RGI_AMR_analysis.md file. In summary, all unique ARGs were identified across all the isolates, then each unique ARG was searched for the number of times it appeared in the isolates (only counting the first copy in the event that multiple copies of the same ARG were present in the same isolate), and then a percentage across all isolates was calculated.

### MDR decision tree models

To investigate key genes involved in multi-drug resistance (MDR), additional data from BV-BRC were downloaded to build machine-learning models, consisting of 352 *E. coli* genomes with corresponding MIC values to at least two different antibiotics. The genome IDs can be found in E coli genomes MDR models.csv. J48 decision tree models were created using *E. coli* genomes in WEKA (71) for 55 pairwise comparisons between 16 antibiotics (Table S3). We opted to use decision trees over more complex models, such as Random Forest and XGBoost, because we needed the interpretability of a single tree so we could screen each gene to merge with our Coinfinder pangenome analysis. Even though there were a possible 240 pairwise comparisons, data only existed for 55. These models represent how MDR arises (we used pairs of antibiotics; therefore, the models can only predict MDR for two antibiotics). The genomes were categorised into “Susceptible” and “Resistant” for individual antibiotics using EUCAST breakpoints (64). The model can predict various phenotype combinations for different antibiotics. For example, a pairwise comparison between two antibiotics could have four different combinations: SS, SR, RS, RR (where S: Susceptible and R: Resistant; and the first letter would correspond to the phenotype of the antibiotic A and the second letter of each pair would correspond to the phenotype to antibiotic B).

The data input for WEKA is an Attribute-Relation File Format (ARFF) file. The attributes (i.e., all eggNOG gene families and AMR phenotype) are listed in the ARFF file. The data follows the attributes in the file and is in the format of the copy number of the gene family or absence (0), and the MDR phenotype (either SS, SR, RS, or RR). Each training file had genomes across all four phenotypes, so the model could predict all four phenotypes. A custom Python script Make_MDR_arff_files.py was used to make the training data files to build the MDR decision tree models. The models were trained using the default parameters and to ensure robustness, we performed multiple validation steps: (1) 10-fold cross validation for all machine-learning models (see Table S2), (2) permutation testing (n = 1,000) to confirm statistical significance and (3) independent validation using the held-out dataset of 352 genomes described earlier and which were not included in the initial pangenome analysis. We calculated the precision, recall, and F1 scores for susceptible and resistant predictions separately and averaged the values. This ensured the entire confusion matrix was considered when we evaluated the models’ statistics.

### Combining pangenomic and decision tree results

We investigated the genes involved in particular MDR phenotypes within the decision trees that are significantly associated or dissociated with each other. To achieve this, we searched for each gene family involved in the 55 MDR decision trees in the Coinfinder output files: “*pairs.tsv” for evidence of association and disassociation. This provided a list of gene families involved in significant associations or disassociation between at least one gene family in the MDR decision trees. To focus this list on significant occurrences between gene families only in the decision tree models, we combined both reduced *pairs.tsv files for associated and dissociated gene families and opened in Cytoscape (70). Gene families in the respective models were labelled as “Tree” and “NA” if not in a model. Then, only gene families in the decision tree models were selected. To identify which AMR phenotype (‘SS’, ‘SR’, ‘RS’, and ‘RR’) the gene family was most associated with, a Chi-squared test was applied to the matrix of positive and negative influences the gene had on different phenotypes (please see schematic Figure 4 for reference). Our decision tree models are always binary, representing branch points distinguishing between the phenotypic consequences of having more or fewer of a gene, we counted how many genomes in our models were found to be resistant or susceptible on either side of the branch point associated with each gene in the decision tree. This gave an overview of the “positive” phenotypic associations with a gene (i.e. the phenotypic consequences of having more of this gene) and or the “negative” phenotypic associations with a gene (i.e. the phenotypic consequences of having fewer of this gene). For example, if the presence of a gene is always associated with resistance, then we should observe in the branch associated with having more of this gene, a larger number of genomes with the resistance phenotype than the susceptible phenotype. Meanwhile, in the branch associated with having fewer (or zero) copies of the gene, we should see no particular association with resistant or susceptible genomes, and its absence cannot be selected against. This would leave a tell-tale difference in the ratios of S:R genomes in the negative branch compared to the positive branch, which could be identified with a simple Chi-squared, Fisher’s exact test, and a *post hoc* test (to confirm the phenotype with which the gene was associated) were performed on each gene matrix for each gene family in each model. This also holds for genes whose presence is associated with susceptibility, allowing the identification of genes important to each phenotype. In the case of where there are 4 phenotypes (SS, SR, RS and RR), the same principle holds, and a *post hoc* test can identify which of the phenotypes is most associated with the more (positive) or fewer (negative) copies of the gene. See Figure 4 for a detailed example. The matrix files were calculated using apply decision tree (see code availability). We then merged the positive and negative interaction matrices into one file using a custom Python script merge_presence_absence_matrix.py (Table S9). The overall matrix file was then read into R to perform the stats (this can be found in Gene_phenotype_association.R). Then, we selected pairs of gene families which were found to be significantly associated (P value = *<* 0.05) with a particular MDR phenotype and compared them to the Coinfinder significantly associated gene network and dissociated gene networks. Lists of gene families present in each decision tree model were created. Then, using a custom Python script (match_DT_to_coinfinder.py), we extracted gene family matches in the Coinfinder network (in which the source and target nodes in the Coinfinder network corresponded to gene families in one decision tree). We then searched for the significantly associated/dissociated gene pairs that we found in *E. coli* in the *P. aeruginosa* Coinfinder networks.

**Figure 4.**
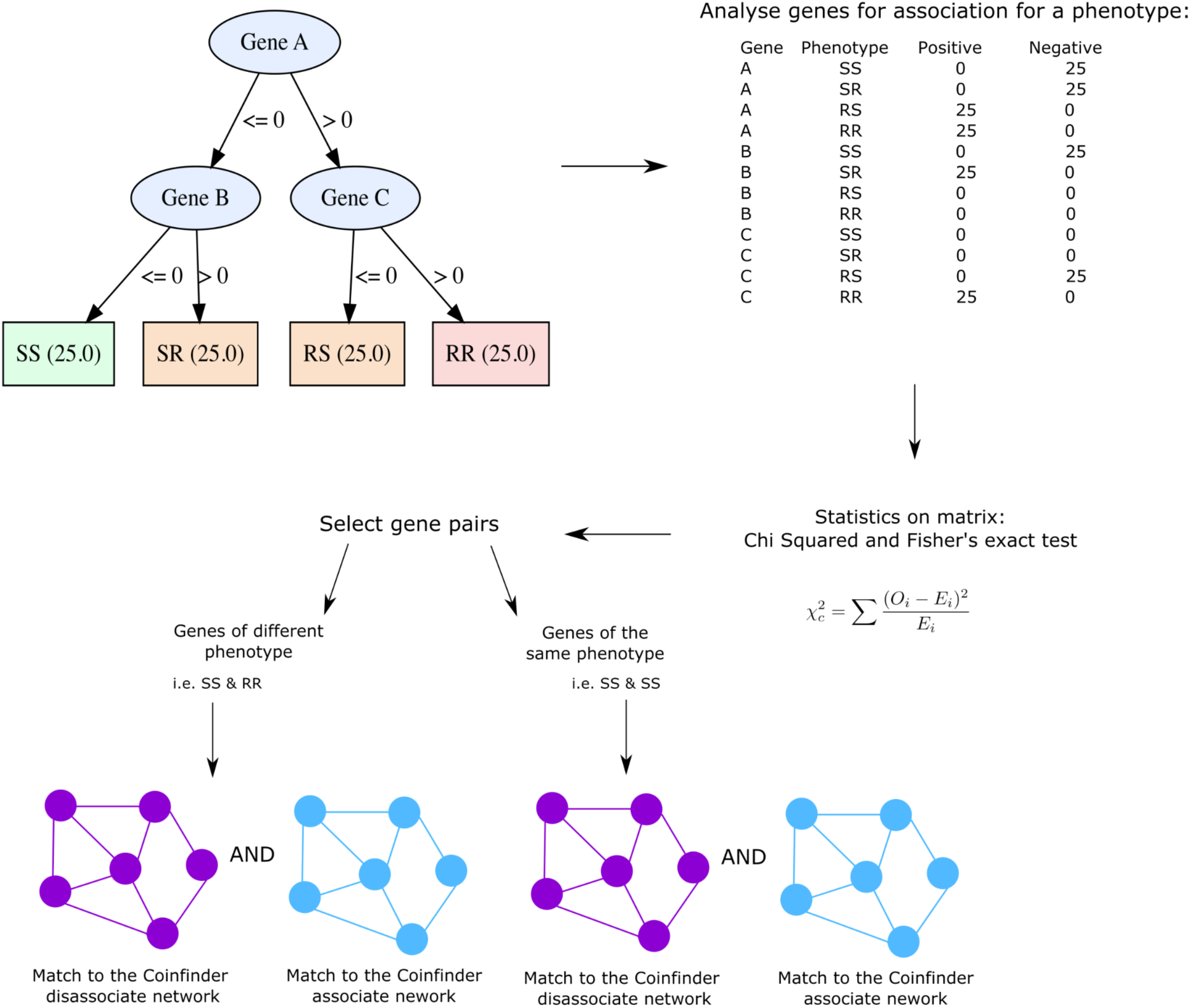
Schematic representation of how gene families within the decision trees are mapped to the Coinfinder networks. The top left shows a representative decision tree, the gene families are analysed in a matrix by assessing how many genomes it is associated with (top right). The matrix for each gene family within each model is then analysed by using a Chi-squared and Fisher’s exact test. Gene families which were found to be significantly associated with a particular AMR phenotype within the decision tree models were then searched in the Coinfinder associated/dissociated networks. The gene families were searched as a pair from one MDR decision tree model in the Coinfinder associated/dissociated networks. See Table S4 for gene families matched to the Coinfinder.

## Acknowledgements

We acknowledge PhD funding from the Department for the Economy, Northern Ireland, to L Dillon. CJ Creevey wishes to acknowledge funding from UKRI MR/Y015223/1 and EU via Horizon 2020 (818368 MASTER and 101000213 Holoruminant).

## Conflict of Interest

No conflict of interest to declare by all authors.

## Author contributions

L Dillon: conceptualization, software, formal analysis, investigation, visualization, methodology, and writing—original draft, review, and editing.

JO McInerney: conceptualization, methodology, and writing, review, and editing.

CJ Creevey: conceptualization, data curation, software, supervision, investigation, visualization, methodology, and writing—original draft, review, and editing.

## Abbreviations

AMR: Antimicrobial resistance
ARFF: Attribute-Relation File Format
ARG: Antimicrobial resistance gene
ATP: Adenosine triphosphate
BLAST: Basic local alignment search tool
BV-BRC: Bacterial and Viral Bioinformatics Resource Center
CARD: Comprehensive Antibiotic Resistance Database
EUCAST: European Committee on Antimicrobial Susceptibility Testing
MDR: Multidrug resistance
MIC: Minimum inhibitory concentration
OSF: Open science framework
PCR: Polymerase chain reaction
PTS: Phosphotransferase system
RGI: Resistance gene identifier
WGS: Whole genome sequence

## Notes

https://osf.io/95k6z/?view_only=b46e8760a93b4e70a849fe4d2210165f

https://github.com/LucyDillon/MDR_pangenome

### Competing Interest Statement

The authors have declared no competing interest.

### Summary of Updates

This updated version has a more focused narrative than the previous version.

