## Supplementary Figure for "Dynamic genomic constraints reveal fitness trade-offs underlying bacterial resistance evolution"

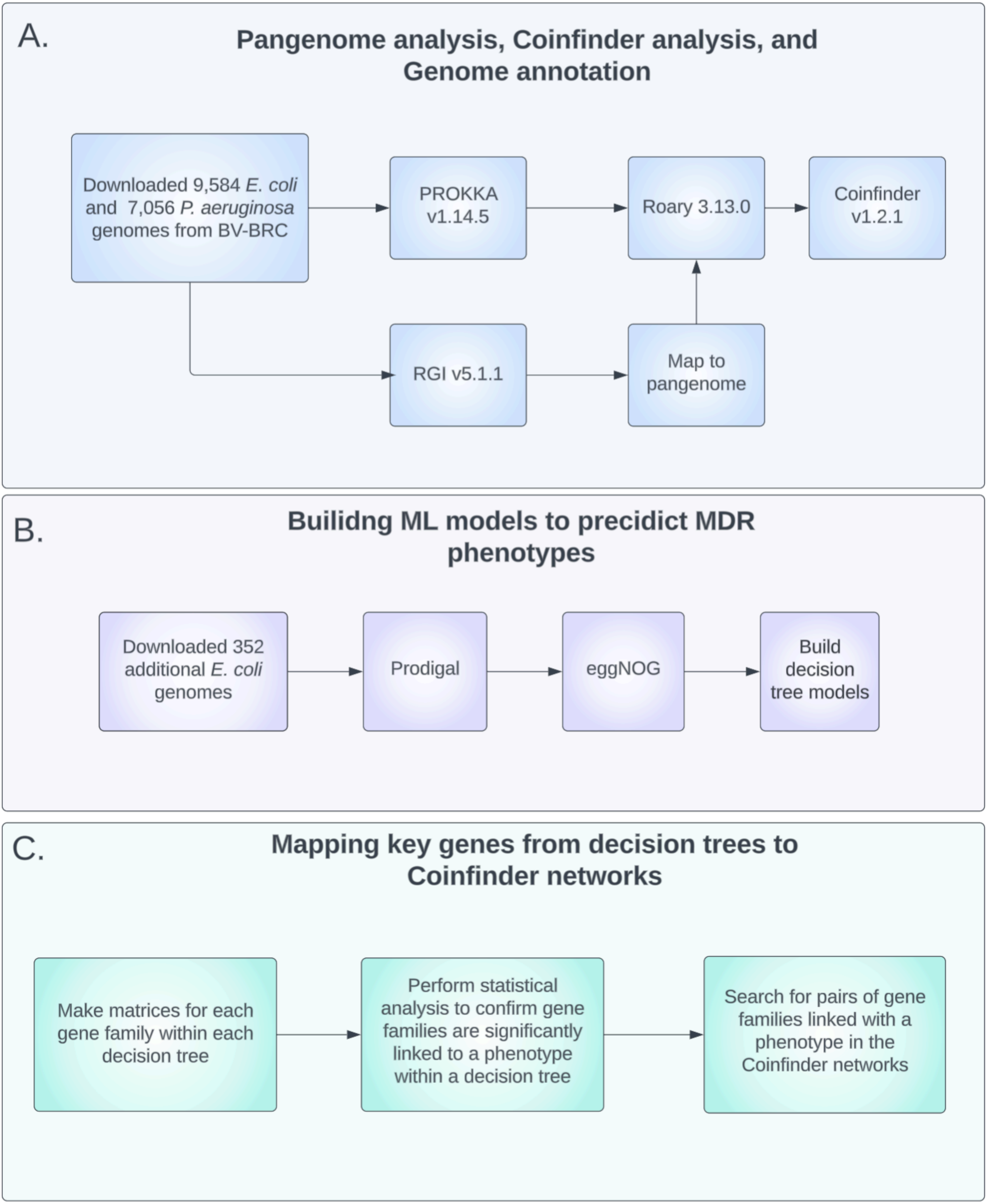
A.
Pangenome analysis, Coinfinder analysis, and Genome annotation
Downloaded 9,584 E. coli and 7,056 P. aeruginosa genomes from BV-BRC
PROKKA v1.14.5
Roary 3.13.0
Coinfinder v1.2.1
RGI v5.1.1
Map to pangenome
B.
Builidng ML models to precidict MDR phenotypes
Downloaded 352 additional E. coli genomes
Prodigal
eggNOG
Build decision tree models
C.
Mapping key genes from decision trees to Coinfinder networks
Make matrices for each gene family within each decision tree
Perform statistical analysis to confirm gene families are significantly linked to a phenotype within a decision tree
Search for pairs of gene families linked with a phenotype in the Coinfinder networks

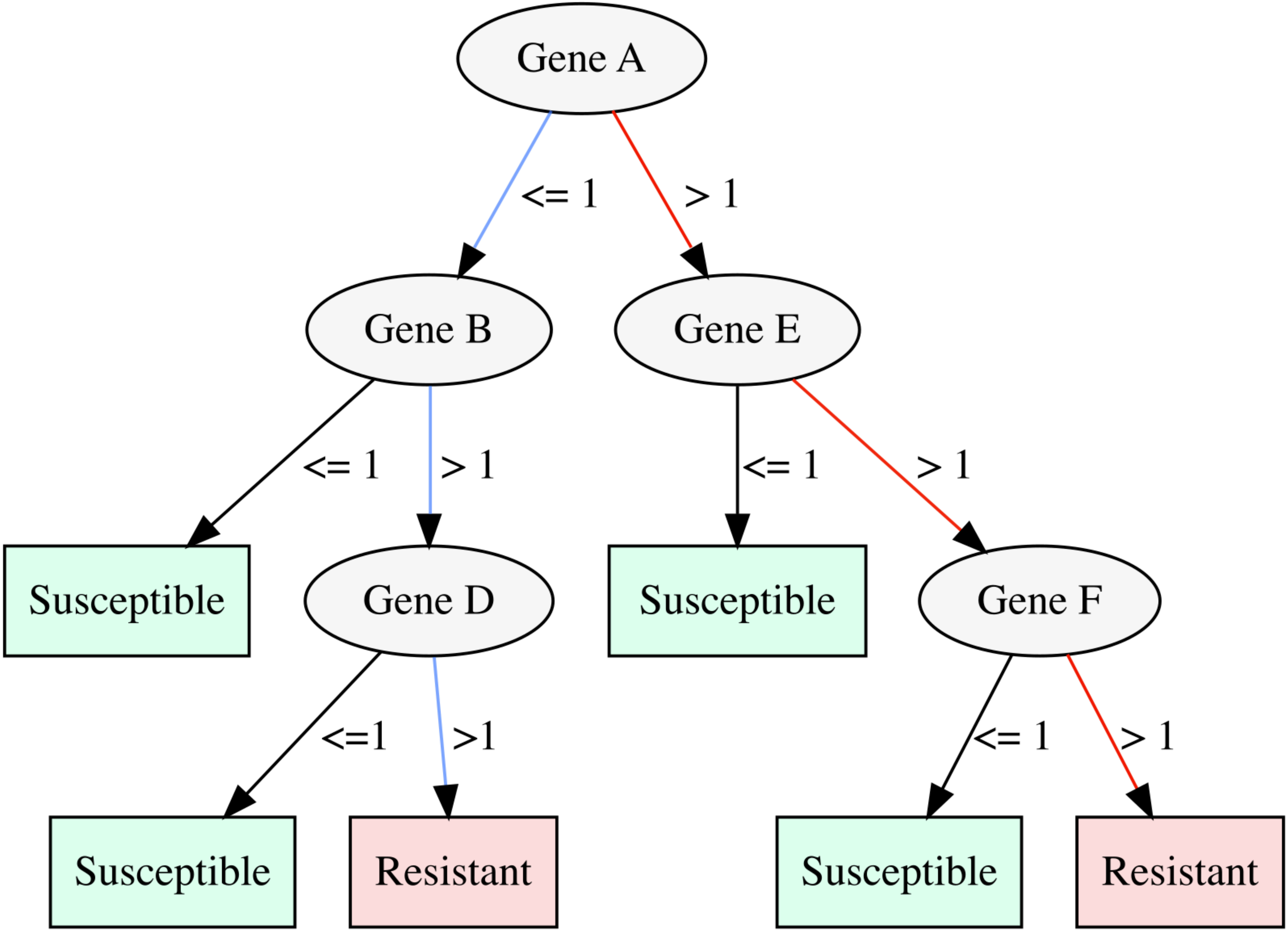
Gene A
<= 1
> 1
Gene B
Gene E
<= 1
> 1
<= 1
> 1
Susceptible
Gene D
Susceptible
Gene F
<=1
>1
<= 1
> 1
Susceptible
Resistant
Susceptible
Resistant

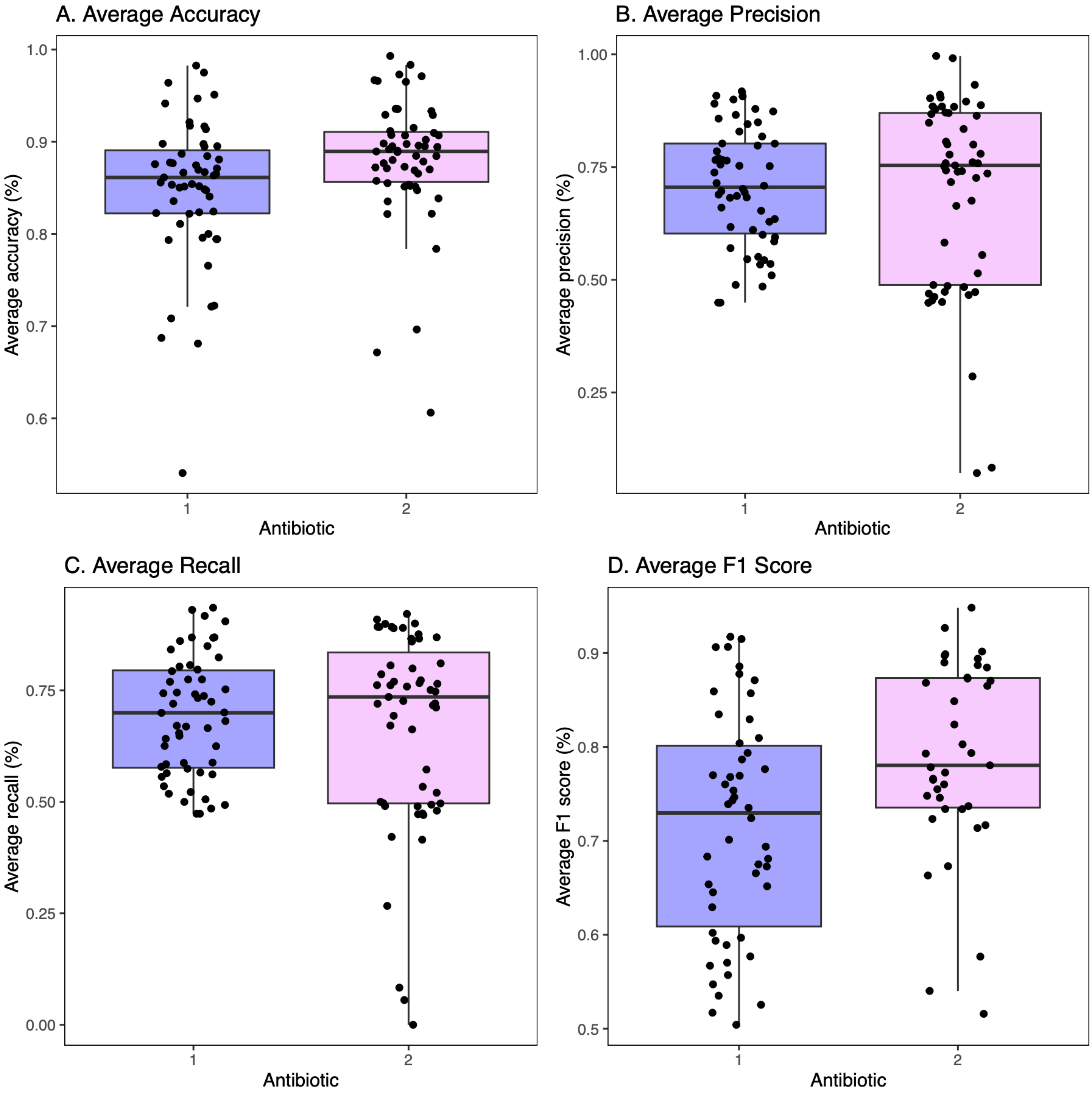
A. Average Accuracy
Average accuracy (%)
1.0
0.9
0.8
0.7
0.6
1
2
Antibiotic
B. Average Precision
Average precision (%)
1.00
0.75
0.50
0.25
1
2
Antibiotic
C. Average Recall
Average recall (%)
0.75
0.50
0.25
0.00
1
2
Antibiotic
D. Average F1 Score
Average F1 score (%)
0.9
0.8
0.7
0.6
0.5
1
2
Antibiotic

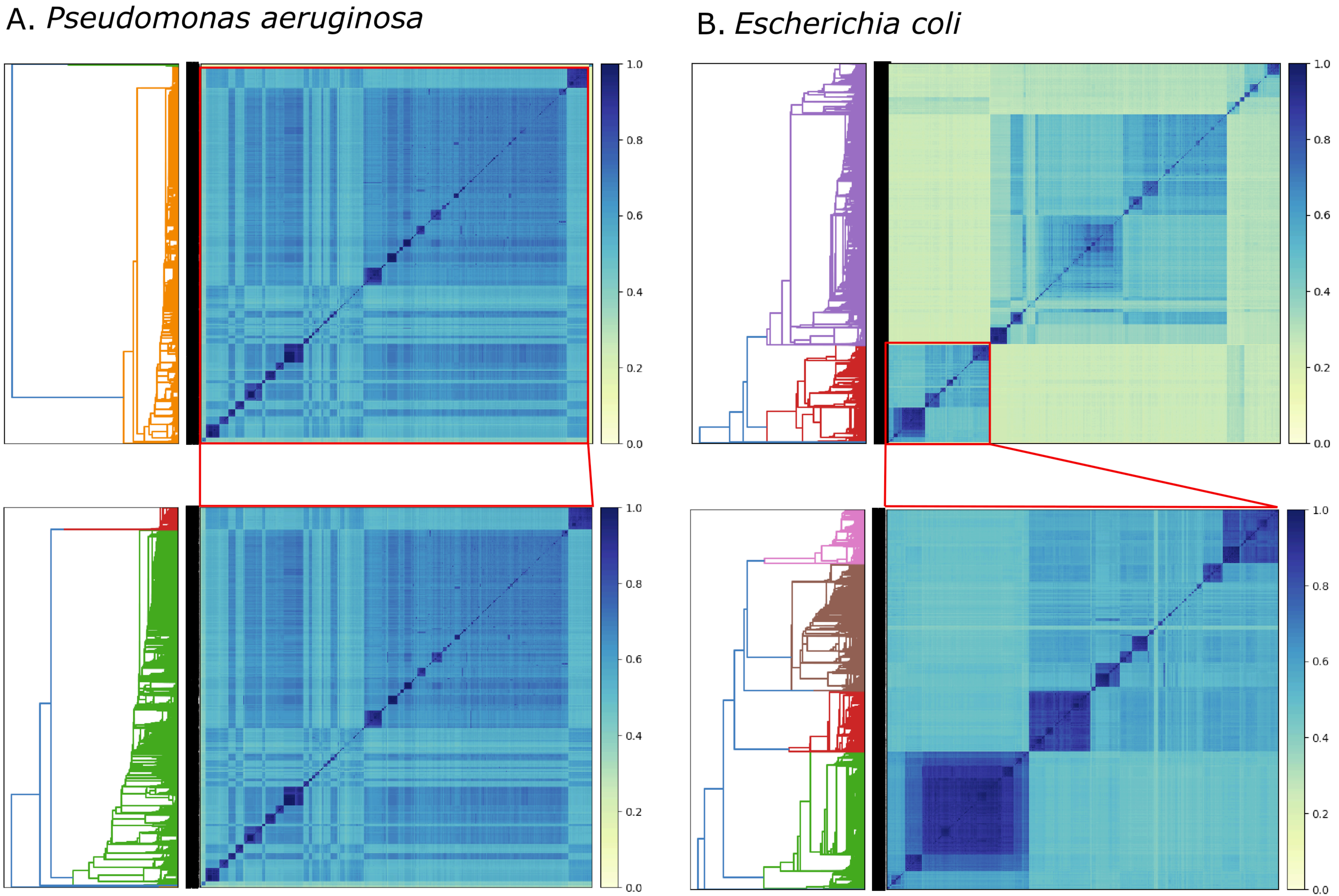
A. Pseudomonas aeruginosa
B. Escherichia coli
1.0
0.8
0.6
0.4
0.2
0.0

Supplementary figure legends:

Figure S1: A workflow of methods used in this study. Part A shows the pangenome analysis, Coinfinder analysis of the pangenome and the genome functional annotation. Part B shows how we built the machine learning models used in this study. Part C shows how we mapped genes significantly associated with a phenotype in a decision tree to the Coinfinder networks.

Figure S2: Example decision tree. To use the decision tree, the input enters the top node (Gene A). Then, according to the presence/absence of the genes, the user will follow the route according to the values defined on the edges. This shows that there are two routes to resistance (red and blue)

Figure S3: A boxplot displaying how the decision tree models performed for the individual antibiotics involved in the dual antibiotic resistance models. Showing statistics accuracy (A), precision (B), recall (C), and F1 score (D). Each data point represents an individual MDR model generated.

Figure S4: Sourmash of *P. aeruginosa* (A) and *E. coli* (B). The heatmaps show the Sourmash results of all genomes against all, using the Jaccard index to cluster the genomes (the higher the Jaccard index, the closer the relation). The y-axis shows the clustering of the genomes, the darker the colour of the heatmap the more closely related. The top two heat maps in A and B are all the genomes sourced from BV-BRC, and the heat maps below represent the final genomes used in the analysis. The reduction in the genomes in parts A and B is due to removing misclassified species, redundancy of genomes, and a subcluster of *E. coli* was chosen to have closer related genomes and reduce the number of genomes (from >37,000 to <10,000).
